# Disruption of augmin-mediated microtubule nucleation in neural stem cells causes p53-dependent apoptosis and aborts brain development

**DOI:** 10.1101/2020.11.18.388694

**Authors:** Ricardo Viais, Sadanori Watanabe, Marina Villamor, Lluís Palenzuela, Cristina Lacasa, Marcos Fariña, Jens Lüders

## Abstract

Microtubules that assemble the mitotic spindle are generated by three different mechanisms: centrosomal nucleation, chromatin-mediated nucleation, and nucleation from the surface of other microtubules mediated by the augmin complex. Impairment of centrosomal nucleation in apical progenitors of the developing mouse brain induces p53-dependent apoptosis and causes non-lethal microcephaly. Whether disruption of non-centrosomal nucleation has similar effects is unclear. Here we show, using mouse embryos, that conditional knockout of the augmin subunit *Haus6* in apical progenitors led to spindle defects and mitotic delay. This triggered massive apoptosis and complete abortion of brain development. Co-deletion of p53 rescued cell death, but brain development was still aborted. This could be explained by exacerbated mitotic errors and resulting chromosomal defects including increased DNA damage. Surviving progenitors had lost apico-basal polarity and failed to organize a pseudostratified epithelium. Thus, in contrast to the centrosomal nucleation pathway, augmin is crucial for apical progenitor mitosis, and, even in the absence of p53, for progression of brain development.

## Introduction

Spindle assembly crucially depends on microtubule nucleation by the γ-tubulin ring complex (γTuRC). During mitosis γTuRC generates microtubules through three different pathways: centrosomal nucleation, chromatin-mediated nucleation, and nucleation from the surface of other microtubules [1-3]. The latter mechanism is mediated by the augmin complex and has been referred to as a microtubule amplification mechanism [4-7]. Augmin binds to the lattice of microtubules generated by the centrosome- and chromatin-dependent pathways and, through recruitment of γTuRC, promotes nucleation of additional microtubules that grow as branches from these sites [8-11]. The existence of multiple nucleation pathways may provide some level of redundancy to spindle assembly, but concerted action by multiple nucleation mechanisms has also been described [1,12]. While functional studies in *Xenopus* egg extract and cultured cell models have generated a wealth of information regarding the types of spindle defects that occur when specific nucleation pathways are compromised, how these defects impinge on cell fate and development remains poorly defined.

Gene mutations that cause functional or numerical centrosome aberrations are associated with primary microcephaly, a developmental disorder that results in reduced thickness of the cerebral cortex. Depletion of cortical progenitors following abnormal mitoses has been identified as a pathogenic mechanism [13-15]. Apical progenitors of the developing cerebral cortex are highly polarized cells. Their cell bodies are positioned in the ventricular zone (VZ), while their apical and basal processes contact the ventricular surface and basal lamina, respectively [16-18]. Prior to mitosis the nucleus migrates apically and mitotic chromosome segregation occurs near the apical surface. Early during cortical development apical progenitors divide symmetrically, expanding the progenitor pool. At later stages they switch to self-renewing asymmetric mitoses, producing a neuron or intermediate progenitor in each division. Centrosomal microtubules were proposed to be at the core of these fate decisions, by controlling the distribution of cell fate determinants through correct positioning of the mitotic spindle[19-21]. Recent work showed that progenitor fate is strongly impacted by mitotic duration. Mitotic delay results in more neurogenic divisions and an increased percentage of progenitors undergoing p53-dependent apoptosis, depleting the progenitor pool [22,23]. Consistently, mitotic delay, premature differentiation and apoptosis have all been observed for centrosome defects in mouse models of primary microcephaly [24-28]. Interestingly, in cases where it has been tested, such as *Cenpj*-or *Cep63*-deficient mice, the reduced cortical thickness was fully rescued by co-deletion of p53, identifying p53-dependent apoptotic cell death as main driver of microcephaly in these models [25,26].

The roles of chromatin-mediated nucleation and augmin-dependent amplification in this context are less clear. Mouse embryos deficient for *Tpx2*, a spindle assembly factor that functions in chromatin-mediated nucleation, abort development after a few rounds of highly abnormal mitotic divisions [29]. Similar observations were made for mouse embryos lacking expression of the augmin subunit *Haus6* [30]. However, since early mouse development occurs in the absence of centrosomes [31], the embryos in the above studies lacked two of the three mitotic nucleation pathways.

Early functional studies by augmin knockdown in cell lines described mitotic defects that ranged from relatively mild for Drosophila cells [4,32] to more severe for human cells [6], suggesting cell type-or organism-specific differences. Consistent with this, the knockout of augmin in *Aspergillus* has no obvious phenotype [33], *Drosophila* augmin mutants are viable with mild mitotic defects observed in only some cell types [32,34], and a zebrafish mutant is also viable but displays defects in the expansion and maintenance of the hematopoietic stem cell pool [35]. A more recent inducible knockout of the augmin subunit *HAUS8* in non-transformed human RPE1 cells caused mild spindle defects before cells underwent p53-dependent G1 arrest, but co-deletion of p53 exacerbated the mitotic phenotype [36]. This response may involve the USP28-53BP1-p53-p21-dependent mitotic surveillance pathway, which is triggered by centrosome loss or prolonged mitosis [37-39], but this was not directly tested.

To uncover the specific role of augmin-mediated microtubule amplification in mitotic spindle assembly and cell fate determination, we sought to study augmin deficiency in centrosome-containing cells *in vivo*. To this end we conditionally knocked out *Haus6* in proliferating apical progenitors in the embryonic mouse brain using nestin promotor-driven Cre expression. We found that augmin is essential for brain development, promoting mitotic progression and preventing p53-dependent progenitor apoptosis. Intriguingly, while absence of p53 promoted growth in *Haus6* knockout brains, this was accompanied by exacerbated mitotic errors and disruption of tissue integrity. Our results show that contrary to centrosomal microtubule nucleation, the augmin-dependent pathway is essential for apical progenitor mitotic progression and survival, and thus for brain development.

## Results

### Augmin is essential for proper development of the mouse forebrain

Previous work has shown that the augmin complex is composed of eight subunits and that depletion of any subunit interferes with augmin assembly and function [4,6,7]. Mouse embryos that completely lack expression of the augmin subunit *Haus6* do not survive the blastocyst stage [30]. In order to test the specific requirement for augmin in proliferating neural progenitors, we generated floxed *Haus6* mice in which exon 1 of the *Haus6* gene is flanked by loxP sequences. We then crossed these mice with mice expressing Cre recombinase under control of the Nestin promoter, to induce *Haus6* knockout in apical progenitors starting around day E10.5 (Fig. 1a; Fig. S1a) [40,41]. In contrast to the full knockout, *Haus6* conditional knockout (*Haus6* cKO) mice passed through all developmental stages and at E13.5 we observed efficient deletion of *Haus6* in the brain (Fig. S1b). Whereas mice with a heterozygous *Haus6* deletion developed normally and were fertile*, homozygous Haus6 cKO* mice died around birth. Analysis of *Haus6* cKO animals at E17.5 showed severe defects in brain development, whereas overall body development appeared normal (Fig. 1b, c; Fig. S1c). Histopathology analysis revealed a strong disruption or absence of different forebrain structures (cortex, thalamus, hypothalamus) and of the cerebellum (Fig. 1c; Fig. S1c). To evaluate whether this was due to agenesis or tissue loss during development, we analyzed embryos at E13.5. Even at this earlier stage brains in *Haus6* cKO embryos displayed severe defects compared to control embryos. Lateral cortexes in *Haus6* cKO embryos were almost completely absent and thalamus structures, while partially formed, displayed a strong reduction in radial thickness (Fig. 1d). Moreover, spaces between tissue structures were filled with cellular debris. These data suggest that, in *Haus6* cKO brains at early developmental stages, formation of structures that would give rise to the cortex, thalamus and hypothalamus is initiated but not completed, leading to tissue loss and abortion of brain development at later stages.

**Figure 1:**
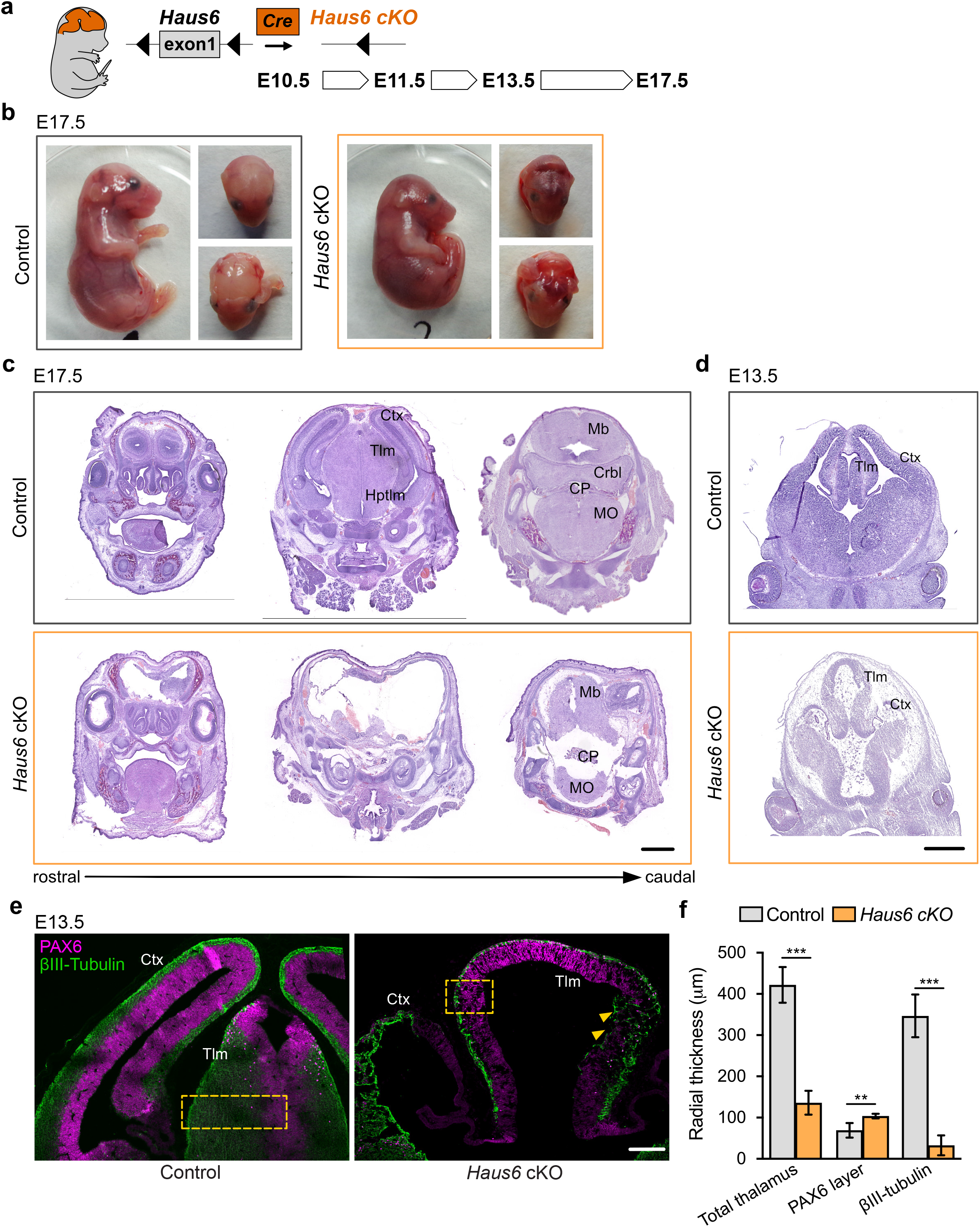
Loss of *Haus6* aborts forebrain development. **(a)** Schematic representation of the experimental strategy used to evaluate the role of augmin during mouse brain development through generation of brain-specific Nestin-Cre *Haus6* cKO embryos. **(b)** Pictures of E17.5 control (*Haus6^fl/fl^* Nestin-Cre^-^) and *Haus6* cKO (*Haus6^fl/fl^* Nestin-Cre^+^) embryos. **(c, d)** Coronal histological sections from (**c**) E17.5 and (**d**) E13.5 control and *Haus6* cKO stained with haematoxylin-eosin. Different brain structures are labeled: Ctx (cortex), Tlm (thalamus), Hptlm (hypothalamus), Mb (midbrain), Crbl (cerebellum), MO (medulla oblongata) and CP (choroid plexus). **(e)** Representative images of the cortex (Ctx) and thalamus (Tlm) of E13.5 control (*Haus6*^fl/wt^ Nestin-Cre^+^) and *Haus6* cKO (*Haus6*^fl/fl^ Nestin-Cre^+^) embryos. Coronal sections were stained against PAX6 (magenta - apical progenitors) and βIII-tubulin (green - neurons). Yellow arrowheads highlight regions of the thalamus where tissue disruption is observed in *Haus6* cKO embryos. Yellow boxes indicate the regions used for quantifications in (**f**). **(f)** Quantification of the total radial thickness of the thalamus in E13.5 embryos and of layers formed by PAX6- and βIII-tubulin-positive cells. n=3 control and n=5 *Haus6* cKO embryos. Plotted values are means, error bars show SD. **P<0.01, ***P<0.001 by two-tailed t-test. Scale bars: (**c**) 1 mm, (**d**) 0.5 mm, (e) 150 μm.

### Loss of augmin impairs mitotic progression in cortical and thalamic neural progenitors

To analyze defective brain development in *Haus6* cKO animals at E13.5 at the cellular level we focused on the thalamus, which was at least partially preserved. We co-stained brain sections with antibodies against PAX6 and βIII-tubulin to label apical progenitors and neurons, respectively. In *Haus6* cKO embryos we observed that the reduced radial thickness in the thalamus was due to a striking thinning of the neuronal layer by ~90% when compared to controls (Fig. 1e, f), indicating severely impaired neurogenesis. In some parts, where tissue organization appeared to be disrupted, we also observed neurons that were misplaced in apical regions (Fig. 1e). To directly test if augmin deficiency impaired mitoses, we identified and quantified mitotic cells in the thalamus using Ser10-phospho-Histone H3 staining. In *Haus6* cKO embryos we observed a ~4-fold increase in the number of mitotic cells in the region closest to the ventricular surface compared to controls, whereas there were no significant differences in more basal regions (Fig. 2a, b). The percentage of *Haus6* cKO mitotic cells in prometaphase was strongly increased, whereas metaphases and ana/telophases were reduced relative to controls (Fig. 2c). Together these observations suggest that augmin deficiency in progenitors of the thalamus leads to a defect in progression to metaphase, causing mitotic delay.

**Figure 2:**
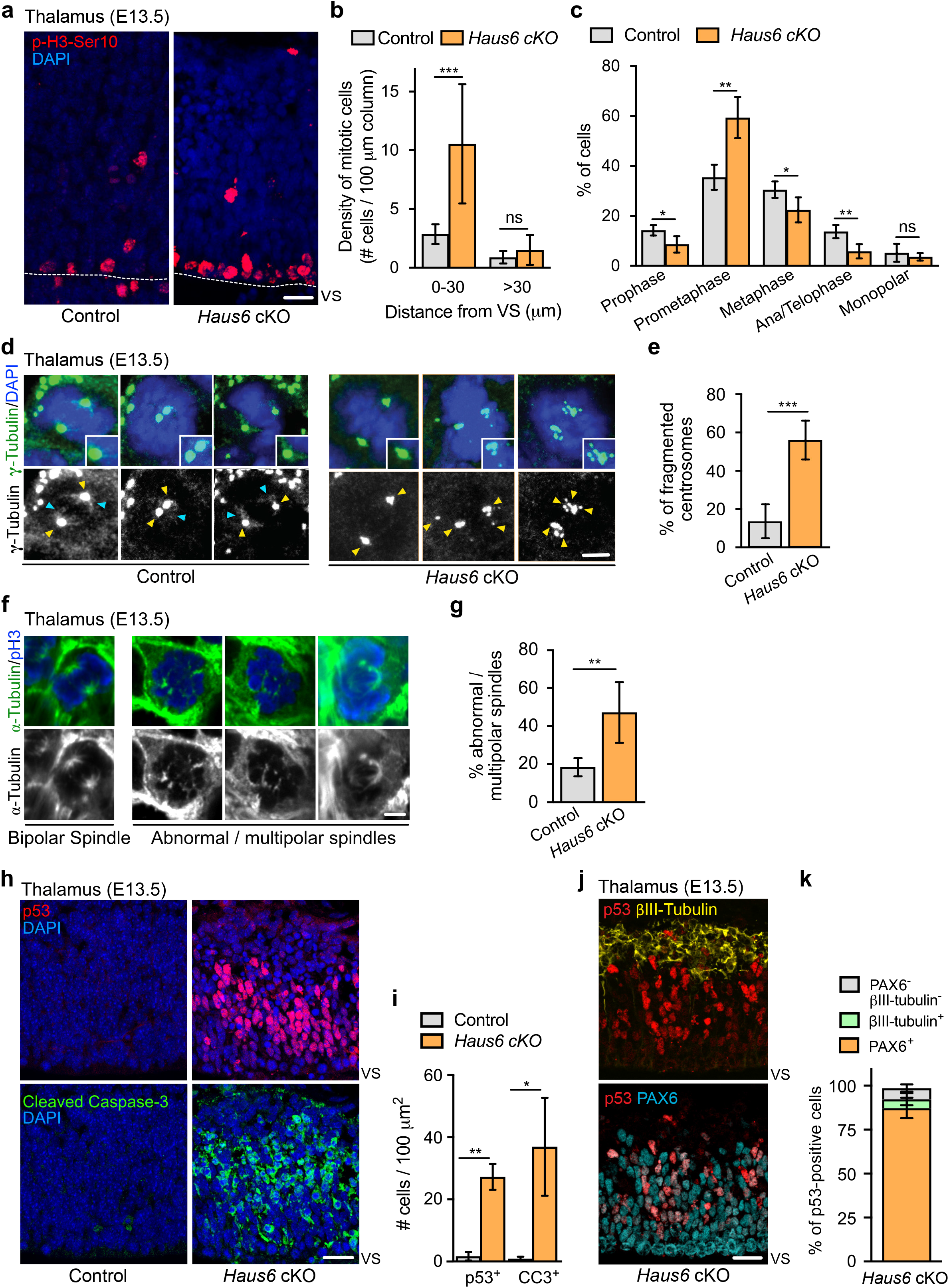
Augmin deficiency in neural progenitors impairs mitotic spindle assembly and induces p53 expression and apoptosis. **(a)** Representative images of phospho-Histone H3 (pH3) positive mitotic cells in the thalamus of E13.5 control (*Haus6*^fl/wt^ Nestin-Cre^+^) and *Haus6* cKO (*Haus6^fl/fl^* Nestin-Cre^+^) embryos. Staining of pH3 in red and DAPI in blue. **(b)** Quantification of the density of mitotic cells close to the ventricular surface (VS) (<30 μm away) and in outer layers of the cortical plate (>30 μm away). n=4 control and n=4 *Haus6* cKO embryos. **(c)** Quantification of mitotic progenitors at the different mitotic stages. n=4 control and n=5 *Haus6* cKO embryos. **(d, f)** Representative images of E13.5 control and *Haus6* cKO mitotic progenitors close to the VS of the thalamus. **(d)** Coronal brain sections were stained with antibodies against γ-tubulin (green) and DAPI to label DNA (blue). Yellow arrowheads point to γ-tubulin staining at spindle poles. Light blue arrowheads point to spindle-associated γ-tubulin staining. **(e)** Quantification of the percentage of mitotic cells in (d) with fragmented centrosomes. **(f)** Coronal sections were stained with antibodies against α-tubulin (green) and pH3 (blue). **(g)** Quantification of the percentage of mitotic progenitors from control and *Haus6* cKO E13.5 embryos displaying abnormal/multipolar spindles. n=5 Control and n=3 *Haus6* cKO embryos. **(h)** Representative images of control (*Haus6*^fl/wt^ Nestin-Cre^+^) and *Haus6* cKO (*Haus6*^fl/fl^ Nestin-Cre^+^) coronal brain sections showing the thalamic region stained with an antibody against p53 (red – upper panel) and the apoptotic marker cleaved caspase-3 (green – lower panel). DNA is labeled with DAPI (blue). **(i)** Quantification of the density of p53 and cleaved caspase-3 positive cells in the E13.5 thalamus in brain sections as shown in (h). n=3 control and n=3 *Haus6* cKO embryos for quantifications of p53 positive cells and n=4 control and n=3 *Haus6* cKO embryos for cleaved caspase-3-positive cells. **(j)** Representative images of *Haus6* cKO coronal brain sections showing the thalamus stained for p53 (red), PAX6 (cyan) and βIII-tubulin (yellow). **(k)** Quantification of the percentage of *Haus6* cKO cells showing induction of p53 and co-express PAX6 (orange), βIII-tubulin (light green) or none of these markers (grey). n=3 E13.5 thalamus regions of different *Haus6* cKO embryos. **(b, c, e, g, i, k)** Plotted values are means, error bars show SD. *P<0.05, **P<0.01, ***P<0.001 by two-tailed t-test. Scale bars: (a) 20 μm, (d, f) 3 μm, (h, j) 25 μm.

To analyze cortical progenitors and since there were no intact cortical structures in *Haus6* cKO brains at E13.5, we analyzed embryos at E11.5. At this stage cortical structures were present suggesting that, as for the thalamus, cortical tissue is originally formed but lost at later stages. Similar to the situation in the thalamus at E13.5, in *Haus6* cKO cortexes at E11.5 the percentage of mitotic progenitors was increased when compared to controls and this occurred specifically in the apical region and not in more basal layers. Again this increase in mitotic cells was due to accumulation in prometaphase (Fig. S2a, b, c, d). Together the data show that augmin plays an important role in allowing timely mitotic progression of apical progenitors in different regions of the developing mouse brain.

To test if augmin-deficient progenitors displayed spindle defects we analyzed brain sections with antibodies against γ-tubulin and α-tubulin (Fig. 2d, e, f, g). Mitotic apical progenitors in the thalamus of control animals displayed strong, centrosomal staining of γ-tubulin at spindle poles and more diffuse γ-tubulin signals along spindle microtubules. In *Haus6* cKO embryos γ-tubulin could not be detected on spindle microtubules. Moreover, in ~56% of cells the staining of γ-tubulin at spindle poles was dispersed into multiple smaller foci, consistent with pole fragmentation (Fig. 2d, e). Indeed, labeling of microtubules by α-tubulin antibodies revealed abnormal spindle configurations, frequently with multiple poles, in about half of the mitotic progenitors in *Haus6* cKO animals (Fig. 2f, g). However, bipolar configurations including at ana/telophase were also observed and cell divisions occurred in *Haus6* cKO progenitors, suggesting that in some cases multipolarity may be avoided through pole clustering and that mitosis was not completely blocked.

Considering that augmin-deficiency caused pole fragmentation, we wondered whether this affected spindle positioning. We measured spindle angles relative to the ventricular surface in dividing apical progenitors in the thalamus and in the cortex of E13.5 and E11.5 *Haus6* cKO embryos, respectively. We found that in both cases the majority of spindles axes were oriented horizontally similar to spindles in control cells (Fig. S2e, f, g). This is consistent with results from previous work showing that the presence of multiple spindle poles in progenitors due to extra centrosomes does not significantly affect spindle orientation [42].

In summary, augmin deficiency in apical progenitors disrupts recruitment of γ-tubulin to spindle microtubules, causes pole fragmentation, and interferes with bipolar spindle assembly and mitotic progression.

### Loss of augmin in neural progenitors induces p53 expression and apoptosis

We sought to determine the fate of progenitors undergoing abnormal mitoses after loss of augmin. We probed thalamus and cortex of E13.5 and E11.5 *Haus6* cKO embryos, respectively, for p53 induction and the presence of the apoptotic marker cleaved caspase-3. Indeed, p53 and cleaved caspase-3 were strongly upregulated in both brain regions (Fig. 2h, i; Fig. S2h), whereas cells positive for these markers were barely found in the corresponding tissues of control embryos. To reveal the identity of cells overexpressing p53, we performed a triple staining with antibodies against p53, the neuronal marker βIII-tubulin and the apical progenitor marker PAX6 (Fig. 2j). This experiment showed that in the *Haus6* cKO thalamus ~87% of the p53-positive cells were also positive for PAX6 and only a minor fraction (~5%) for βIII-tubulin (Fig. 2k). Moreover, we observed that PAX6-positive progenitors displaying p53 induction were exclusively interphase cells, based on the presence of intact nuclei. We concluded that p53 induction occurred specifically in augmin-deficient progenitors, after exit from abnormal mitoses.

Together the data suggests that mitotic spindle defects in *Haus6* cKO progenitors are not catastrophic *per se*, but efficiently trigger apoptotic cell death upon completion of mitosis.

### Co-deletion of p53 in *Haus6* cKO embryos rescues apoptosis but not forebrain development

Since massive apoptosis in *Haus6* cKO brains was correlated with p53 induction, we wondered whether cell death was p53-dependent and the cause of aborted brain development. To address this, we crossed *Haus6* cKO mice with p53 KO mice (Fig. 3a). Strikingly, at E13.5, a stage at which *Haus6* cKO brains displayed massive apoptosis, lacked cortical structures and had a poorly developed thalamus, *Haus6* cKO p53 KO brains showed only minimal signs of apoptosis and there was some growth in the regions where cortex and thalamus would be expected to form (Fig. 3b, c, d). This growth was enhanced when compared to the single *Haus6* cKO brains, but seemed to lack the layered organization observed in control brains at this stage (Fig. 3b). At E17.5, however, when thalamus and cortex were well formed in controls, in *Haus6* cKO p53 KO embryos cortex and thalamus structures appeared thin and undeveloped (Fig. 3e). Moreover, as observed for *Haus6* cKO embryos, *Haus6* cKO p53 KO animals were not viable and died around birth.

**Figure 3:**
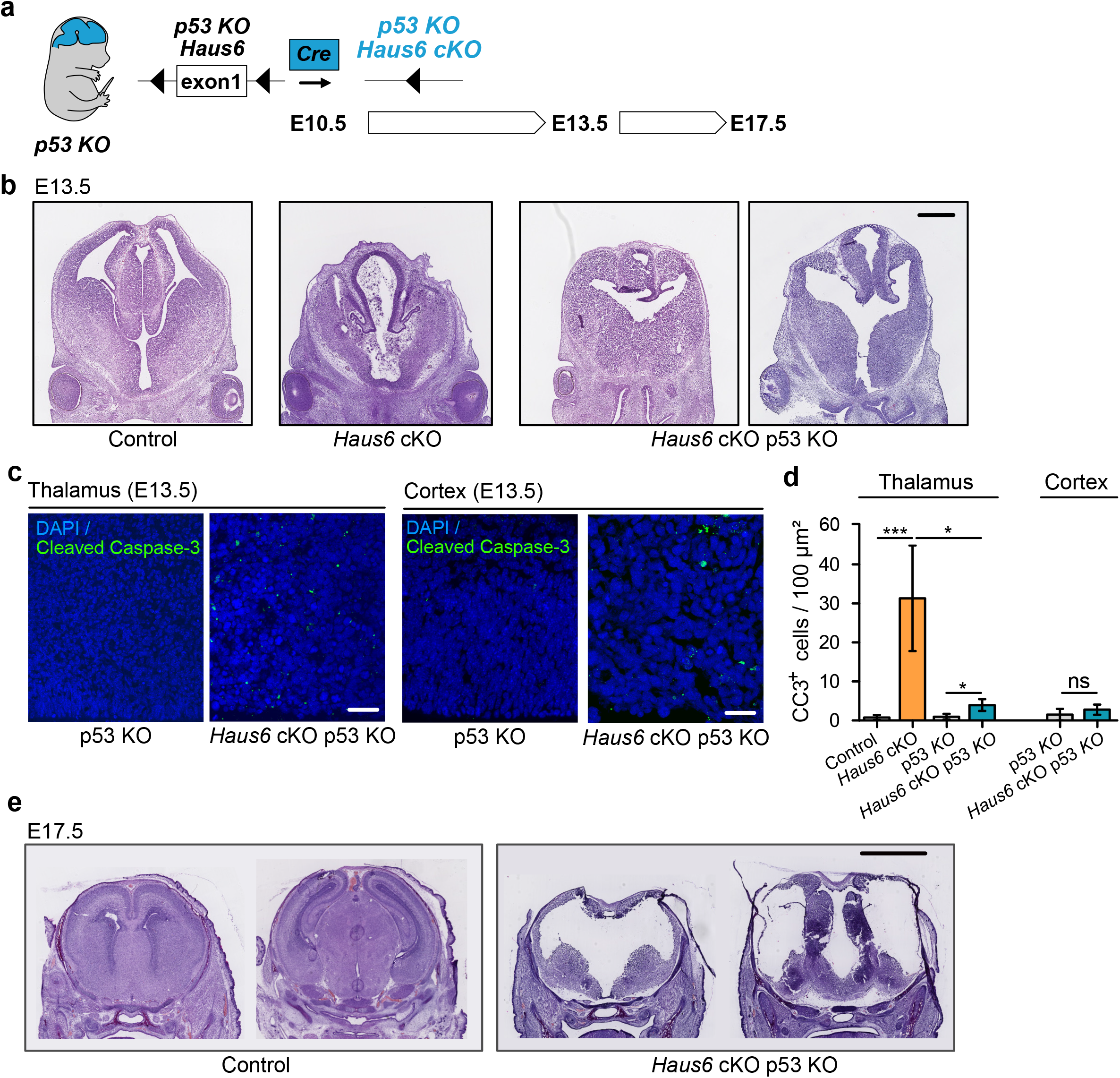
Co-deletion of p53 rescues apoptosis but not abortion of forebrain development. **(a)** Schematic overview showing the experimental strategy used to generate *Haus6* cKO p53 KO embryos, to test p53 dependency of the brain development phenotypes observed in *Haus6* cKO embryos. **(b)** Coronal sections of E13.5 control (*Haus6^fl/wt^* Nestin-Cre^+^), *Haus6* cKO (*Haus6*^fl/fl^ Nestin-Cre^+^) and *Haus6* cKO p53 KO (*Haus6*^fl/fl^ Nestin-Cre^+^ *p53*-^/-^) embryos stained with haematoxylin-eosin. **(c)** Coronal sections of the thalamus and cortex of E13.5 p53 KO Control and *Haus6* cKO embryos stained against the apoptotic marker cleaved caspase-3 (green). DNA was labeled by DAPI (blue). **(d)** Quantification of the density of cleaved caspase-3 positive cells in the E13.5 thalamus and cortex in brain sections as shown in (c). n=5 control, n=5 Haus6 cKO, n=4 p53 KO control and n=3 *Haus6* cKO p53 KO embryos for quantifications in the thalamus and n=4 p53 KO control and n=4 Haus6 cKO p53 KO embryos for quantifications in the cortex. Plotted values are means, error bars show SD. *P<0.05, **P<0.01, ***P<0.001 by two-tailed t-test. **(e)** Coronal sections of E17.5 Control and *Haus6* cKO p53 KO embryos stained with haematoxylin-eosin. Scale bars: (b) 0.5 mm; (c) 40 μm, 25 μm (cortex); (e) 2 mm.

In summary, massive apoptosis in *Haus6* cKO brains is rescued in *Haus6* cKO p53 KO brains, promoting growth in the affected brain regions, but this growth is not productive for proper brain development.

### Loss of p53 exacerbates mitotic defects caused by augmin deficiency

Next we examined how co-deletion of *Haus6* and *p53* affected mitosis in proliferating progenitors. Similar to *Haus6* cKO alone, *Haus6* cKO p53 KO embryos also had an increased density of mitotic cells in the cortex and the thalamus as revealed by Ser10-phospho-Histone H3 staining (Fig. 4a, b, c, d). Closer examination of the mitotic figures in E13.5 cortexes revealed severe mitotic defects in *Haus6* cKO p53 KO embryos. In addition to defective bipolar spindle assembly in prometaphase and metaphase cells, we also observed various abnormalities in post-metaphase cells. Compared to p53 KO control littermates there was a strong increase in the number of defective anaphases and telophases including multipolar spindle configurations, lagging chromosomes and micronuclei formation (Fig. 4e, f, g). We also noticed that a fraction of *Haus6* cKO progenitors displayed enlarged nuclei in interphase, which may be indicative of aneuploidy/polyploidy (Fig. 4h). Considering that there were very few apoptotic cells in the double KO brains (Fig. 3c), we wondered how continued proliferation would affect mitoses in progenitors. We compared mitotic defects in *Haus6* cKO and in *Haus6* cKO p53 KO embryos at E13.5 in the thalamus, a structure that was present in embryos of both genotypes at this stage. We found that mitotic defects were more severe in the *Haus6* cKO p53 KO brains when compared to *Haus6* cKO brains (Fig. 4i, j).

**Figure 4:**
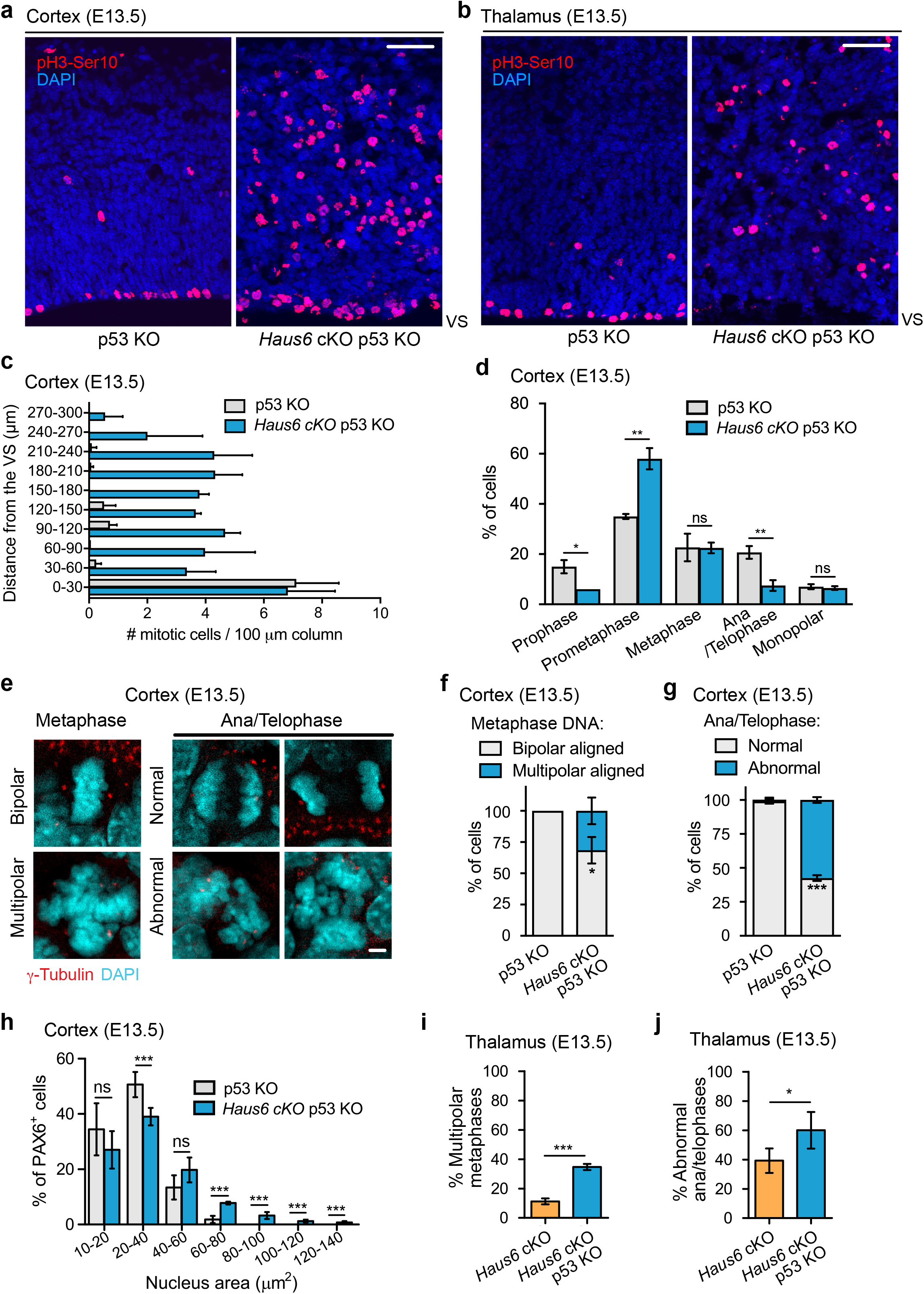
Co-deletion of p53 exacerbates mitotic defects caused by augmin deficiency. **(a, b)** Representative coronal sections of the **(a)** cortex and the **(b)** thalamus of E13.5 p53 KO control (*Haus6^fl/wt^* Nestin-Cre^-^ p53^-/-^) and *Haus6* cKO p53 KO embryos (*Haus6^fl/fl^* Nestin-Cre^+^ p53^-/-^). Sections were stained with phosphorylated Histone H3 antibody (red – mitotic cells) and DAPI to stain DNA (blue). **(c)** Quantification of the density of progenitors undergoing mitosis in the cortex at the indicated distances in μm from the ventricular surface (VS). **(d)** Quantification of mitotic progenitors at the indicated mitotic stages. (**e**) Examples of normal, bipolar mitotic stages and of stages with multipolar and other abnormal configurations in the cortex of E13.5 control and *Haus6* cKO p53 KO embryos, respectively. Coronal sections were stained with an antibody against γ-tubulin to label spindle poles (red) and DAPI (cyan) to label DNA. **(f)** Quantification of the percentage of metaphase cells displaying aligned chromosomes with bipolar (white) and multipolar (blue) configuration in the cortex of embryos with the indicated genotypes. **(g)** Quantification of the percentage of normal and abnormal ana/telophases in the cortex of embryos with the indicated genotypes. **(c, e, f, g)** n=3 p53 KO control and n=2 *Haus6* cKO p53 KO embryos. Plots show mean values, error bars indicate SD. **(h)** Quantification of the nucleus area in interphase PAX6-positive progenitors in the cortex of E13.5 p53 KO control and *Haus6* cKO p53 KO embryos. n=5 p53 KO control and n=4 *Haus6* cKO p53 KO embryos. **(i)** Quantification of the percentage of metaphase cells displaying aligned chromosomes with multipolar configuration in the thalamus of embryos with the indicated genotypes. **(j)** Quantification of the percentage of abnormal ana/telophases in the thalamus of embryos with the indicated genotypes. **(i, j)** n=4 p53 KO control and n=3 *Haus6* cKO p53 KO embryos. **(c,d,f,g,h,i,j)** Plots show mean values, error bars indicate SD. *P<0.05, **P<0.01, ***P<0.001 by two-tailed t-test. Scale bars: (a, b) 50 μm (e) 5 μm.

### Mitotic errors in augmin-deficient progenitors correlate with DNA damage

Since mitotic errors can cause DNA breaks [43], we probed brain tissue of *Haus6* cKO p53 KO embryos for the presence of the DNA-repair marker γH2AX. Indeed, at E13.5 the percentage of cells with interphase nuclei displaying DNA damage was strongly increased in both the cortex and thalamus when compared to controls (Fig. 5a, b, c, d). Side-by-side comparison of γH2AX staining in E13.5 thalamus of *Haus6* cKO and *Haus6* cKO p53 KO embryos showed that augmin deficiency led to increased DNA damage relative to controls and that absence of p53 further increased this effect (Fig. 5c, d). Thus, the extent of mitotic defects that we observed in *Haus6* cKO and *Haus6* cKO p53 KO embryos was correlated with a concomitant increase in DNA damage.

**Figure 5:**
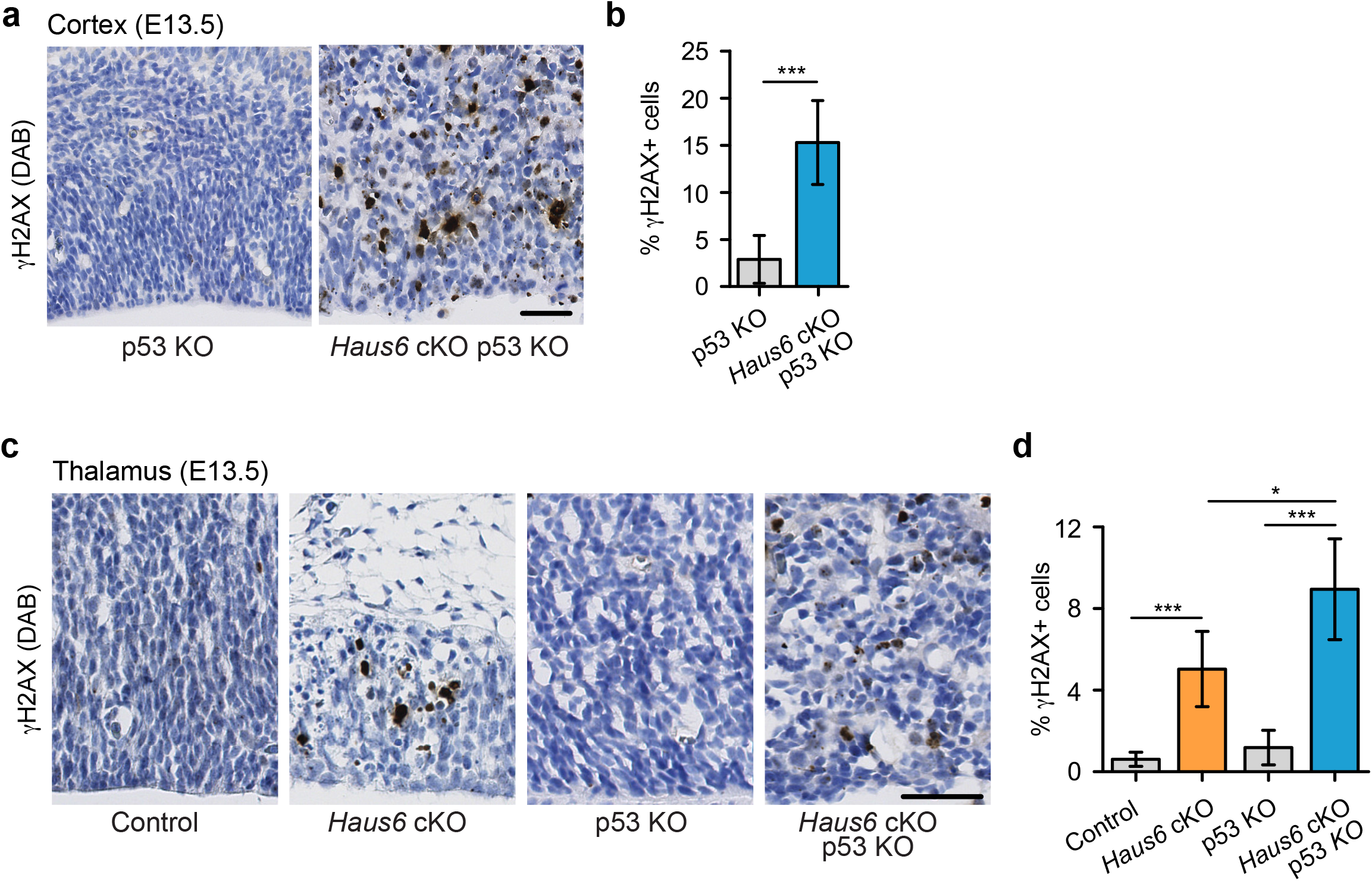
Impaired mitosis in augmin-deficient progenitors causes DNA damage. **(a)** Representative images of a region of the cortex of E13.5 p53 KO control (*Haus6*^fl/wt^ Nestin-Cre^+^ p53^-/-^) and *Haus6* cKO p53 KO (*Haus6*^fl/fl^ Nestin-Cre+ p53^-/-^) embryos. Coronal sections were stained by IHC with an antibody against γH2AX (brown). **(b)** Quantification of the percentage of cells overexpressing γH2AX in the E13.5 cortex. n=8 sections from 4 embryos for p53 KO control and n=7 sections from 4 embryos for *Haus6* cKO p53 KO. **(c)** Representative images of the region of the thalamus of E13.5 control (*Haus6*^fl/wt^ Nestin-Cre^+^ p53^+/+^), *Haus6* cKO (*Haus6^fl/fl^* Nestin-Cre^+^ p53^+/+^), p53 KO control and *Haus6* cKO p53 KO embryos. **(d)** n=6 sections from 3 embryos for control, n=8 sections from 4 embryos for *Haus6* cKO, n=8 sections from 4 embryos for p53 KO and n=6 sections from 3 embryos for *Haus6* cKO p53 KO genotypes. **(b,c)** Plots show mean values, error bars indicate SD. *P<0.05, **P<0.01, ***P<0.001 by two-tailed t-test. Scale bars: (a,c) 45 μm.

### Proliferation in the absence of augmin disrupts neuroepithelium integrity

Apart from the aberrant mitoses in *Haus6* cKO p53 KO progenitors, the distribution of mitotic figures within the tissue was also highly abnormal. Whereas in control and *Haus6* cKO brains the vast majority of mitotic figures with condensed chromosomes were observed in the apical region, near the ventricular surface (Fig. 2a, b; Fig. S2a, c), in *Haus6* cKO p53 KO brains most of the mitotic figures were distributed throughout the tissue including more basal regions (Fig. 4a, b, c).

The presence of large numbers of basally positioned mitotic figures in the cortex and thalamus of *Haus6* cKO p53 KO embryos could indicate that apical progenitors had delaminated, that their nuclei did not migrate to the apical region prior to division, or that the cells displaying mitotic defects in basal layers were not apical progenitors. The latter possibility was tested by PAX6 staining (Fig. 6a). Whereas in the cortex of p53 KO controls PAX6-positive cells were confined to the ventricular zone, well separated from more basally positioned neurons labelled by βIII-tubulin staining, in *Haus6* cKO p53 KO cortex PAX6-positive cells localized indiscriminately in basal and apical regions of the cortex, largely overlapping with regions populated by βIII-tubulin-positive neurons (Fig. 6a, d). Interestingly, TBR2-positive intermediate progenitors, residing in the subventricular zone in control sections, had also lost this confined localization in *Haus6* cKO p53 KO cortexes (Fig. 6b, e). During development, apical progenitors maintain a bipolar structure with their centrosomes lining the ventricular surface, a configuration that is readily visualized by γ-tubulin staining in control embryos (Fig. 6c). In *Haus6* cKO p53 KO embryos apical centrosome localization was strongly reduced and sometimes completely lost (Fig. 6c, f). Occasionally, centrosome clusters were observed in subventricular regions, which was never observed in controls (Fig. 6c). Together these observations suggested that progenitors in *Haus6* cKO p53 KO cortexes were not only incorrectly positioned, but had also lost their polar organization. To assess this more directly, we stained for nestin, an intermediate filament protein specifically expressed in apical progenitors. In the cortex of control embryos, nestin-stained progenitors displayed a highly polarized, apico-basal morphology and a laterally aligned arrangement within the tissue (Fig. S3a). In contrast, polarized morphology and lateral alignment were completely disrupted in progenitors of *Haus6* cKO p53 KO embryos (Fig. S3a). Consistent with these observations, staining with α-tubulin antibodies revealed that microtubules displayed apico-basal organization in control cells, running along the length of the highly polarized cell bodies (Fig. S3b, c). In contrast, microtubules in *Haus6* cKO p53 KO progenitors lacked apico-basal orientation and appeared disorganized (Fig. S3b, c).

**Figure 6:**
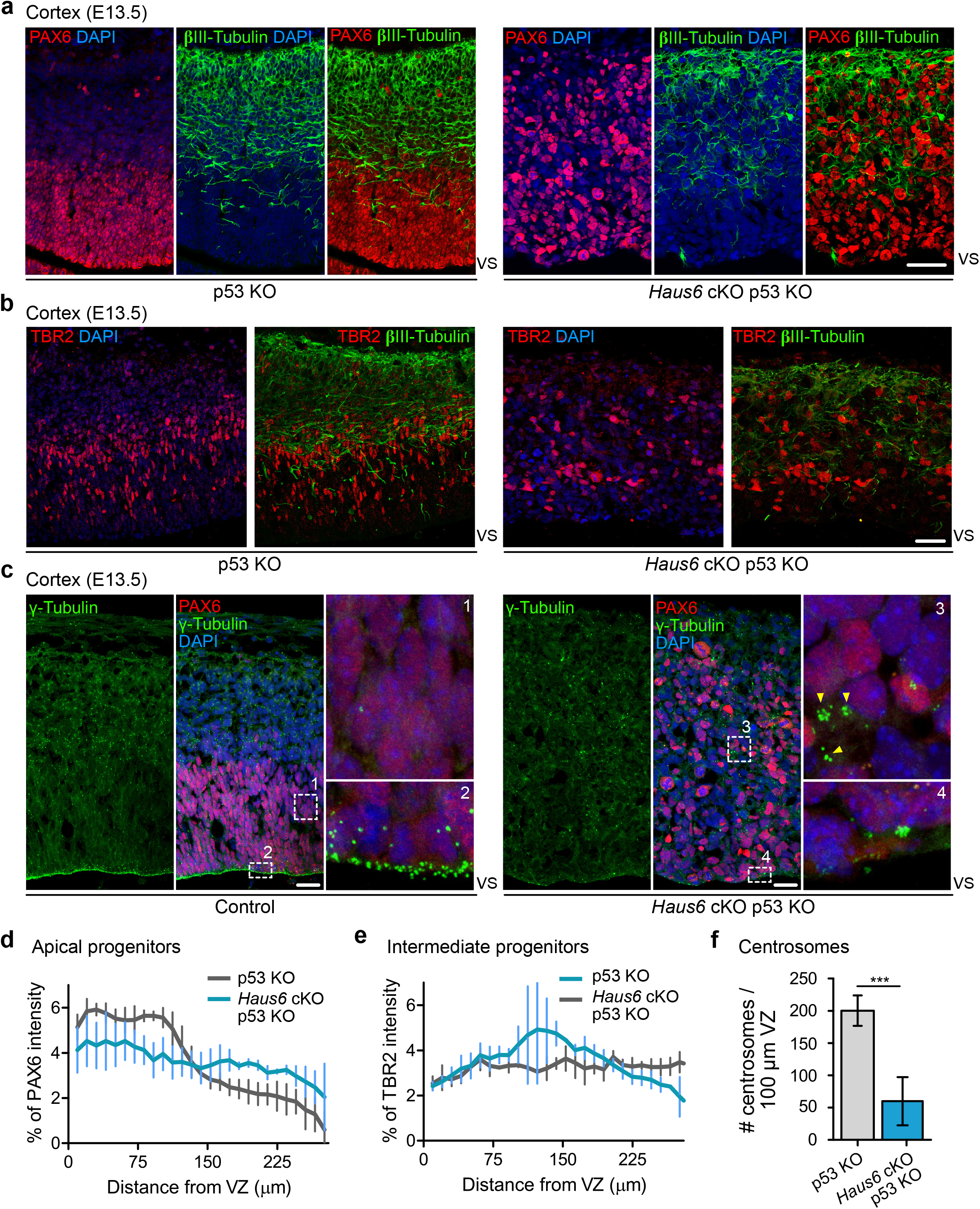
Co-deletion of *Haus6* and p53 leads to loss of cortical layering. **(a, b)** Representative images of the E13.5 cortex from p53 KO control (*Haus6^fl/wt^* Nestin-Cre^+^ p53^-/-^) and *Haus6* cKO p53 KO (*Haus6*^fl/fl^ Nestin-Cre^+^ p53^-/-^) embryos stained with antibodies against PAX6 (a) or TBR2 (b) (red) and the neuronal marker βIII-tubulin (green). DNA was stained with DAPI. **(c)** Representative images of E13.5 cortex from control (*Haus6^fl/fl^* Nestin-Cre^-^ p53^-/-^) and *Haus6* cKO p53 KO (*Haus6^fl/fl^* Nestin-Cre^+^ p53^-/-^) embryos. Coronal sections were co-stained with antibodies against γ-tubulin (green) and PAX6 (red). DNA was stained with DAPI. Magnifications of the boxed regions labeled with 1, 2, 3 and 4 are shown. In the magnified region labeled with 4, yellow arrowheads point to ectopic clusters of centrosomes. **(d, e)** Distribution of PAX6 and TBR2 staining in sections as in (a) and (b), respectively. Intensity values were averaged into 9,8 μm-thick bins and plotted as percentage of total intensity. Lines connect mean values and error bars display SD. (d) n=5 p53 KO and n=4 *Haus6* cKO p53 KO embryos. (e) n=2 p53 KO and n=2 *Haus6* cKO p53 KO embryos. **(f)** Quantification of the density of centrosome number at the ventricular surface of the cortex of E13.5 p53 KO and *Haus6* cKO p53 KO embryos. n=4 p53 KO and n=4 *Haus6* cKO p53 KO embryos. Plots show mean values and error bars display SD. ***P<0.001 by two-tailed t-test. Scale bars: (a, b) 35 μm; (c) 25 μm.

Together these data suggest that in *Haus6* cKO p53 KO embryos apical progenitors had lost their polarized organization and divided ectopically. As a result neuroepithelium integrity was severely disrupted.

## Discussion

The mitotic spindle serves to segregate the replicated chromosomes faithfully into two daughter cells. This task is carried out by spindle microtubules and a multitude of proteins that nucleate, organize and remodel these microtubules during mitotic progression. Here we have analyzed the contribution of one of three different microtubule nucleation pathways, augmin-mediated microtubule amplification, to mitotic spindle assembly in proliferating neural progenitor cells during mouse brain development. Previous work found that impairment of centrosomal microtubule nucleation in apical progenitors slowed mitotic spindle assembly and progression, leading to p53-dependent apoptosis and causing microcephaly [24-28]. Similarly, augmin-deficiency also impaired spindle assembly, delayed mitosis, and induced p53-dependent apoptosis. In agreement with previous functional studies in cell lines, augmin-deficient progenitors displayed fragmented spindle poles, but this did not significantly impair spindle positioning. The most important outcome of these defects was cell death. Our finding that the large majority of cells positive for expression of p53 and the apoptotic marker cleaved caspase-3 were PAX6-positive interphase cells, suggests that cell death occurred after completion of abnormal mitoses. Despite the similarities with centrosome defects, the *Haus6* conditional knockout phenotype is much more severe. Rather than leading to microcephaly, augmin deficiency completely aborted brain development. To our knowledge this has not been reported for any other microtubule regulator affecting mitotic spindle assembly and progression. How can this be explained? While mitotic defects and apoptosis were also observed after loss of centrioles by conditional *CenpJ/Cpap/Sas4* knockout [26] and amplification of centrosome number by PLK4 overexpression [42], the specific spindle defects caused by augmin deficiency may be a more potent trigger of apoptotic cell death than defects resulting from centrosome abnormalities. It should be noted that a more recent *Cenpj* conditional knockout mouse model displayed more severe disruption of forebrain structures, causing lethality a few weeks after birth [27]. Still, these defects seem less severe than what we observed after augmin knockout. One may expect that preventing cell death in augmin-deficient progenitors would, at least to some degree, rescue brain development. Codeletion of p53 in *Haus6* cKO mice largely rescued apoptosis, revealing that cell death was p53-dependent, but did not rescue brain development and lethality. In the absence of apoptosis, augmin-deficient progenitors likely underwent repeated cycles of abnormal mitoses, leading to increasingly severe mitotic abnormalities. This behavior has recently been described after induced knockout of the augmin subunit *HAUS8* in the RPE1 cell line. Whereas *HAUS8* knockout in a p53 wild type background only mildly impaired mitosis before cells arrested in G1, co-deletion of p53 eliminated cell cycle arrest and exacerbated mitotic defects [36]. Consistent with this possibility, *Haus6* cKO p53 KO progenitors had more severe mitotic defects than *Haus6* cKO cells, including lagging chromosomes and multipolar spindles at post-metaphase stages, and displayed increased DNA damage. We have not formally tested whether cell death in augmin-deficient progenitors involves the recently described, USP28-53BP1-p53-p21-dependent mitotic surveillance pathway [44]. However, our results show that during brain development cells that have undergone erroneous mitosis are efficiently eliminated in a p53-dependent manner, and that this occurs independently of whether the cause is centrosomal or non-centrosomal. The pole-fragmentation phenotype in augmin-deficient mitotic progenitors may be comparable to mitoses in the presence of extra centrosomes, as described in mice overexpressing PLK4 [42]. In these animals co-deletion of p53 also exacerbated mitotic defects and aneuploidy, but the outcome was still a microcephalic brain [42]. In contrast, in the case of *Haus6* cKO p53 KO progenitors in our study, continued proliferation was not productive for brain development. While some cortical structures were present at E13.5, they lacked a pseudostratified epithelial organization. *Haus6* cKO progenitors had lost their characteristic, highly polarized morphology and formed a disorganized cell mass that was intermingled with βIII-tubulin-positive differentiated neurons, in both apical and basal regions. Considering that the polarized apical progenitor morphology is integral to the organization of the neuroepithelium, providing scaffold function and guidance for translocating basal progenitors and migrating neurons, it is not surprising that these defects lead to abortion of brain development.

Exacerbated mitotic errors and resulting DNA damage as a result of continued proliferation are a reasonable explanation for the severely disrupted tissue integrity in *Haus6* cKO p53 KO brains. However, we cannot exclude that additional roles of augmin contribute to this phenotype. For example, augmin may promote progenitor polarity by generating and/or maintaining the apico-basal interphase microtubule array. Recent work has shown that experimentally altered spindle positioning in progenitors can lead to loss of apical membrane. This can be compensated for by re-extension of the apical process and re-integration of the apical foot at the ventricular surface [47]. Assuming a role of augmin in progenitor polarity, this process may be impaired in augmin-deficient cells. Consistent with this possibility, microtubules in *Haus6* cKO p53 KO progenitors appeared disorganized, lacking the apico-basal alignment that is observed in control cells. However, it is unclear whether this is cause or consequence of the loss of polarized cell morphology. It should also be noted that augmin nucleates microtubules in post-mitotic neurons, affecting their morphogenesis and their migration [45,46], which could contribute to tissue disruption in *Haus6* cKO p53 KO brains.

In summary, our work shows that, in contrast to centrosomal nucleation, augmin-mediated microtubule amplification in neural apical progenitors is essential for brain development and cannot be compensated for by the chromatin- and centrosome-dependent nucleation pathways. As in the case of progenitors lacking centrosomal nucleation, mitotic delay caused by augmin deficiency triggers p53-dependent apoptosis. While cell death can be prevented by co-deletion of p53, the specific defects that result from the loss of augmin are sufficient to completely abort brain development, independent of p53 status.

## Methods

### Generation and husbandry of mice

Nestin-Cre *Haus6* cKO were obtained by crossing *Haus6* floxed (*Haus6^fl^*) mice with B6.Cg-Tg(Nes-cre)1Kln/J mice. *Haus6* floxed Neo mice (*Haus6*^fl-Neo^) (Acc. No. CDB1218K, http://www2.clst.riken.jp/arg/mutant%20mice%20list.html) were generated as described [30]. To generate *Haus6* floxed mice (*Haus6^fl^*) (RBRC09630, Accession No.CDB1354K (http://www2.clst.riken.jp/arg/mutant%20mice%20list.html), *Haus6*^fl-Neo^ mice were crossed with C57BL/6-Tg(CAG-flpe)36Ito/ItoRbrc (RBRC01834)[48]. The resultant mice without the PGK-neo cassette (*Haus6* flox mice) were maintained by heterozygous crossing(C57BL/6N background). B6-Tg(CAG-FLPe)36 was provided by the RIKEN BRC through the National Bio-Resource Project of the MEXT, Japan. B6.Cg-Tg(Nes-cre)1Kln/J) mice were a gift from Maria Pia Cosma (CRG, Barcelona, Spain) and previously purchased from Jackson Laboratories. To obtain Nestin-Cre *Haus6* cKO *p53* KO mice, mice carrying the floxed *Haus6* (*Haus6*^fl^) and Nestin-Cre alleles were crossed with mice lacking p53. p53-deficient mice were purchased from Jackson Laboratories. All the mouse strains were maintained on a mixed 129/SvEv-C57BL/6 background in strict accordance with the European Community (2010/63/UE) guidelines in the Specific-Pathogen Free (SPF) animal facilities of the Barcelona Science Park (PCB). All protocols were approved by the Animal Care and Use Committee of the PCB/University of Barcelona (IACUC; CEEA-PCB) and by the Departament de Territori I Sostenibilitat of the Generalitat de Catalunya in accordance with applicable legislation (Real Decreto 53/2013). All efforts were made to minimize use and suffering.

### Mice Genotyping

Genotyping was performed by PCR using genomic DNA extracted from tail or ear biopsies. Biopsies were digested with Proteinase-K (0.4 mg/mL in 10 mM Tris-HCl, 20 mM NaCl, 0.2% SDS, 0.5 mM EDTA) overnight at 56°C. DNA was recovered by isopropanol precipitation, washed in 70% ethanol, dried and resuspended in H2O. To detect *Haus6* wt (800 bp), *Haus6* floxed (1080 bp) and *Haus6* KO (530 bp) alleles by PCR the following pair of primers was used: mAug6KO_FW (5’ - CAACCCGAGCAACAGAAACC-3’) and mAug6KO_Rev (5’-CCTCCCACCAACTACAGACC-3’). These PCRs were run for 35 cycles with an annealing temperature of 64.5°C. To detect the transgenic Cre-recombinase allele in Nestin-Cre cKO mice (100bp) primers olMR1084 (5’-GCGGTCTGGCAGTAAAAACTATC-3’) and olMR1085 (5’-GTGAAACAGCATTGCTGTCACTT-3’) were used. For this PCR, primers olMR7338 (5’-CTAGGCCACAGAATTGAAAGATCT-3’) and olMR7339 (5’-GTAGGTGGAAATTCTAGCATCATCC-3’) were used as internal control (324 bp). These PCRs were run for 35 cycles with an annealing temperature of 51.7°C.

### Histology, immunofluorescence and immunohistochemistry

For histopathology analysis of mouse embryos, timed pregnant female mice were euthanized, embryos were removed, and following euthanasia, embryonic heads were fixed in 4% paraformaldehyde in phosphate buffer saline (PBS) pH 7.4 for 48 hours. Following overnight cryoprotection in 30% sucrose in PBS, tissue was embedded in OCT (Tissue-Tek) and frozen in liquid nitrogen-cooled isopentane. For tissue histological analysis, 10 μm thick cryosections were prepared, placed on glass slides and processed for either hematoxilin/eosin staining using standard protocols or for immunofluorescence staining. For immunofluorescence staining, cryosections were thawed at room temperature, washed with PBS and subjected to heat-mediated antigen retrieval in citrate buffer (10 mM citric acid) at pH6, as required. Tissue sections were permeabilized with PBS containing 0.05% TX100 (PBS-T 0.05%) for 15 minutes and blocked with blocking solution (10% goat serum diluted in PBS-T 0.1%). Sections were then incubated overnight at 4°C with primary antibodies diluted in blocking solution. The next day, after washing with PBS-T 0.05%, sections were incubated for 60 minutes with Alexa-Fluor conjugated complementary secondary antibodies and DAPI to stain DNA. Sections were again washed with PBS-T 0.05% and mounted with Prolong Gold antifading reagent (Thermo Fisher). Immunohistochemistry was performed using 7 μm cuts. Prior to immunohistochemistry antigen retrieval was performed using Tris-EDTA buffer pH 9 for 20 min at 97°C using a PT Link (Dako – Agilent). Quenching of endogenous peroxidase was performed by a 10 min incubation with Peroxidase-Blocking Solution (Dako REAL S2023). Blocking was done in M.O.M. blocking reagent (MKB-2213, Vectorlabs), 5 % of goat normal serum (16210064, Thermo Fisher) mixed with 2.5 % BSA diluted in Envision Flex Wash buffer (K800721, Dako - Agilent) and with Casein solution (ref: 760-219, Roche) for 60 and 30 min, respectively. Mouse monoclonal anti-phospho-Histone H2AX (Ser139), clone JBW301 (05-636, Millipore) was diluted 1:500 with EnVision FLEX Antibody Diluent (K800621, Dako, Agilent) and incubated for 120 min. The secondary antibody used was a polyclonal goat anti-Mouse immunoglobulins/HRP (P0447, Dako-Agilent). Antigen-antibody complexes were revealed with 3-3’-diaminobenzidine (K3468, Dako). Sections were counterstained with hematoxylin (Dako, S202084) and mounted with Mounting Medium, Toluene-Free (CS705, Dako) using a Dako CoverStainer. Specificity of staining was confirmed by using a mouse IgG1 isotype control (ab81032, Abcam).

### Antibodies

The following antibodies were used for immunofluorescence in tissue sections: α-tubulin (Sigma, clone DM1A, dilution 1:500), acetylated α-tubulin (Sigma, #T6793, dilution 1:500), β3-tubulin (AbCam #ab18207, dilution 1:1000), β3-tubulin (Biolegend #801201, dilution 1:1000), cleaved caspase-3 (Novus Biologicals #MAB835, dilution 1:500), γ-tubulin (ExBio, clone TU-30, 1:500), Nestin (Cell signaling #4760, dilution 1:300), p53 (Cell signaling #CST2524S, dilution 1:500), PAX6 (Biolegend #901301, dilution 1:300), phosphorylated-Histone3 (Milipore #06-570, dilution 1:1000), TBR2 (Abcam #ab23345, dilution 1:200). Phosphorylated-Histone H2AX (Ser139) (clone JBW301, Milipore #05-636, dilution 1:500) primary antibody was used for immunohistochemistry in brain sections. Secondary antibodies used: Alexa Fluor 488- and 568-conjugated anti-mouse IgG, Alexa Fluor 488-, 568- and 633-conjugated anti-mouse IgG1, Alexa Fluor 488-conjugated anti-mouse IgG2a, Alexa Fluor 488-, 568- and 633-conjugated anti-rabbit IgG (all raised in goat from Life Technologies, dilution 1:500) and HRP-conjugated anti mouse IgG raised in goat (Dako-Agilent #P0447, dilution 1:500).

### Image acquisition and analysis

Histology sections stained with hematoxilin/eosin (Fig. 1c, 1d, 3b, 3e; Fig. S1c) or used for immunohistochemistry (Fig. 5a, 5c) were imaged with the digital slide scanner Nanozoomer 2.0 HT from Hamamatsu and processed with NDP.view 2 software from Hamamatsu. Immunofluorescence labeled histology sections in Fig. 1e, 2a, 2d, 2h, 2j, 3c, 4a, 4b, 6 and Fig. S2, S3 were imaged with a Leica TCS SP5 laser scanning spectral confocal microscope. Confocal Z-stacks were acquired with 0.5 μm or 1 μm of step size depending on the experiment and using laser parameters that avoided the presence of saturated pixels. Immunofluorescence-labeled histology sections shown in Fig. 2f and Fig. 4e were imaged with a Zeiss 880 confocal microscope equipped with an Airyscan. In the images shown in Fig. 2f, for the Superresolution Airyscan mode a 63x magnification, 1.4NA oil-immersion lens with a digital zoom of 1.8x was used. The z-step between the stacks was set at 0.211 μm. In the images shown in Fig. 4e, for the Fast Airyscan mode a 40x magnification 1.2 NA multi-immersion lens with a digital zoom of 1.8x was used. The Z-step between the stacks was set at 0.5 μm. XY resolution was set at 1588×1588. Airyscan raw data were preprocessed with the automatic setting of Zen Black. Additional image processing and maximum intensity z-projections were done in ImageJ software. In each experiment, serial brain sections from multiple animals per genotype were analyzed (details in figure legends).

Radial thickness of the thalamus was measured with ImageJ as the distance between the ventricular surface and the basal surface of this brain region in E13.5 embryos. In the same regions, radial thickness of the area occupied by PAX6 and βIII-tubulin cell populations was measured.

For mitotic density cell counts the thalamic/cortical wall was divided into 30-μm thick bins from the apical to basal surfaces. The number of mitotic phospho-Histone H3 positive cells was counted in each bin and normalized to the column width of the region analyzed. Mitotic density in each bin was expressed as the number of mitotic cells per 100 μm of column. Centrosome integrity in mitotic cells dividing close to the apical surface of the thalamus/cortex was analyzed by quantifying the percentage of cells displaying unfocused/fragmented spindle poles, each composed of multiple γ-tubulin dots. Mitotic spindle integrity was analyzed in cells dividing close to the apical surface of the thalamus and the percentage of cells displaying abnormal, non-bipolar organized spindles was quantified.

To evaluate p53 expression, cell death and expression of the DNA-damage marker phosphorylated Histone H2AX in the embryonic forebrain, representative images of the thalamus/cortex containing the entire apico-basal axis of the tissue were selected. The number of p53 and cleaved caspase-3 positive cells was counted and divided by the area of the selected region. To evaluate the cell population overexpressing p53 in the thalamus, coronal sections were co-stained against p53, PAX6 and βIII-tubulin. Cells where the p53-positive nucleus was co/stained with PAX6 were counted as PAX6-positive. Cells where the p53-positive nucleus didn’t co-stain for PAX6 and was surrounded by a cytoplasmic βIII-tubulin signal were considered as βIII-tubulin-positive. To evaluate the expression of phosphorylated Histone H2AX, image files obtained with the Nanozoomer 2.0 HT slide scanner were opened with the image analysis software QuPath [49]. The number of phosphorylated Histone H2AX-positive cells was divided by the total amount of hematoxylin-stained cells in the specific tissue, counted using the QuPath software. In all experiments, for each brain at least two coronal tissue sections were quantified.

To measure interphase nucleus size in cortical neural progenitors (Fig. 4h), tissue sections were immunostained with PAX6 antibodies and DAPI to label DNA. The area of nuclei in PAX6-positive cells in the cortex was measured in z-stack images using the “Positive cell detection” tool of QuPath software. Mitotic cells were excluded from this analysis.

To quantify the distribution of neural progenitors within the cortex, cryosections providing lateral views of the cortex were immunostained against PAX6 or TBR2. In both cases, the “Plot Analysis” tool of ImageJ was used to measure signal intensity along the apico-basal axis of the cortex. Measurements were grouped into 9.8 μm-wide bins and the average value for each bin was plotted as percentage of the sum of all bin intensities.

For analysis of mitotic spindle orientation, cryosections providing coronal views of the thalamus/cortex were immunostained with DAPI and the mitotic DNA marker phosphorylated-Histone H3 and the centrosome/spindle pole marker γ-tubulin. The orientation of the mitotic spindle was then determined by measuring the angle between the pole-to-pole axis and the ventricular lining.

### Statistics

All graphics with error bars are presented as means with standard deviation. To determine statistical significance between samples, unpaired two-way Student t-test was used. Statistical calculations and generation of graphs were performed in Excel or Graphpad Prism6 (ns = not significant, * P<0.05, ** P<0.01, *** P<0.001).

## Acknowledgements

General: We are grateful to Gohta Goshima (Nagoya University, Japan) for generously providing floxed *Haus6* mice that were generated in his laboratory and for feedback on the manuscript. We acknowledge excellent support by the IRB Barcelona Histopathology and Advanced Digital Microscopy core facilities for help with sample preparation and analysis, and by the Mouse Mutant Core facility for deriving floxed *Haus6* mice from sperm samples. We thank Travis Stracker (IRB Barcelona, Spain) for mouse cage space and discussion, Eduardo Soriano and Antoni Parcerisas (University of Barcelona, Spain) for technical help and discussions, Irina Matos (The Rockefeller University, USA) for helpful comments on the manuscript, and Pia Cosma (CRG, Barcelona, Spain) for providing Nestin-Cre mice.

## Author contributions

R.V. designed experimental strategies, planned and executed animal experiments, ensured compliance with animal welfare regulations, planned and set up mouse breedings, performed stainings and quantifications, prepared figures and helped with manuscript writing. S.W. designed the strategy for generating the conditional *Haus6* knockout and generated floxed *Haus6* mice. M.V. assisted with animal experiments. L.P. assisted with planning and setting up of animal breedings. M.F. assisted in staining and quantification of cryosections. C.L. supervised animal experiment planning and compliance with animal welfare regulations at IRB Barcelona, and assisted in mouse genotyping. J.L. conceived and supervised the study, and wrote the manuscript.

---

**Figure S1:**
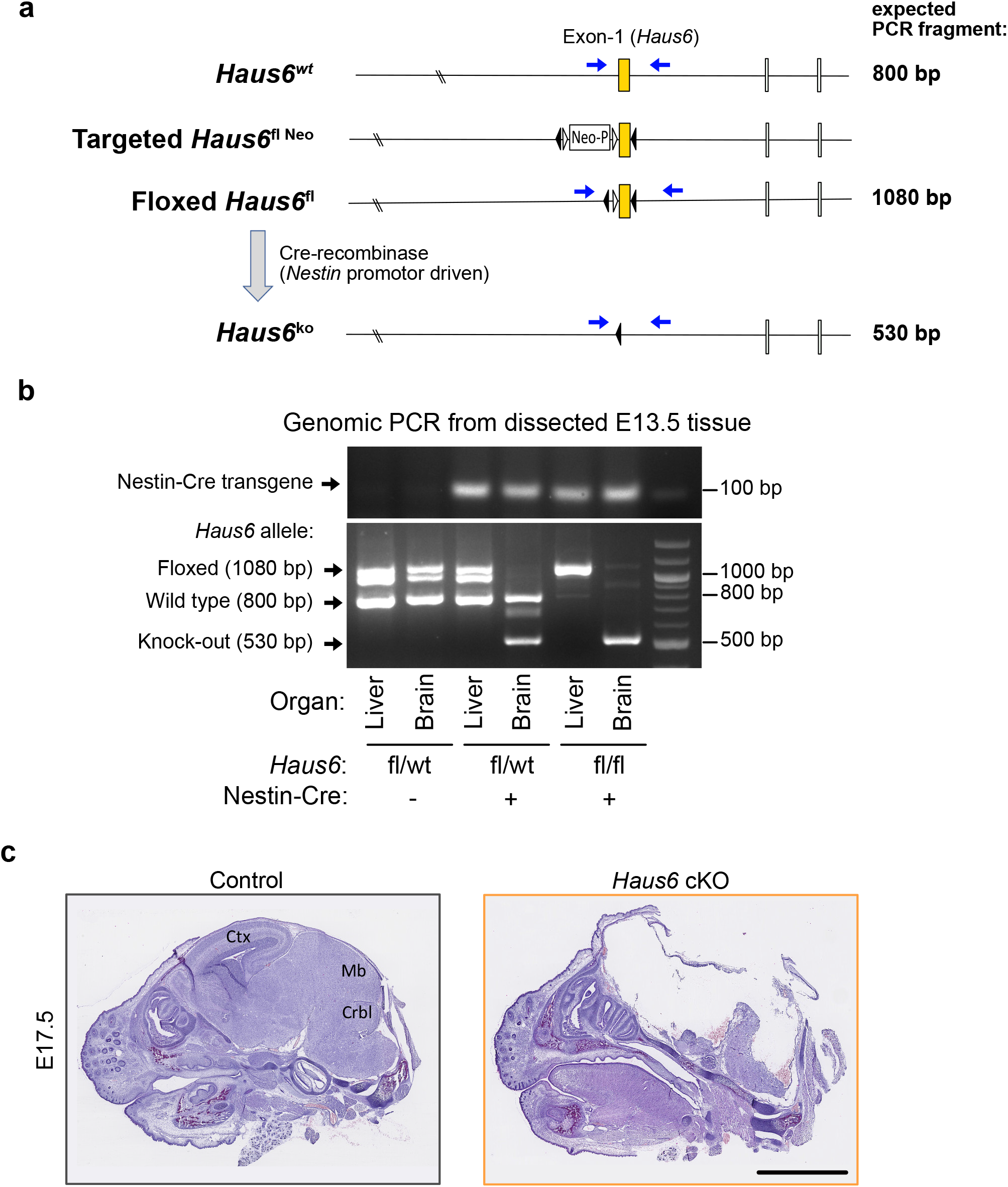
Forebrain structures are absent in *Haus6* cKO embryos at E17.5. **(a)** Schematic representation of the wild type *Haus6* allele (wt), the targeted allele (fl Neo), the floxed *Haus6* allele (fl) and the Haus6 knockout allele (ko). Positions of exons 1–3, loxP sequences (black triangles), flippase recognition target sequences (FRT) (white triangles) and neomycin resistance cassette (Neo) are shown. Positions of PCR primers are indicated by blue arrows and expected PCR fragments are indicated. **(b)** DNA gels showing genomic PCRs from dissected liver and forebrain of E13.5 embryos with different genotypes. In the upper gel, amplification of a 100 bp fragment using Nestin-Cre recombinase primers indicates the presence of the transgene. In the lower gel, amplification of a 1080, 800 and/or 570 bp fragment reveals the presence of *Haus6* floxed, wild type or knockout alleles, respectively, in the indicated genotypes and tissues. Genomic PCR products were run on a 3% agarose gel. **(c)** Sagittal histological sections from E17.5 control (*Haus6*^fl/fl^ Nestin-Cre^-^) and *Haus6* cKO (*Haus6*^fl/fl^ Nestin-Cre^+^) embryos stained with haematoxylin-eosin. Scale bars: (c) 2.5 mm.

**Figure S2:**
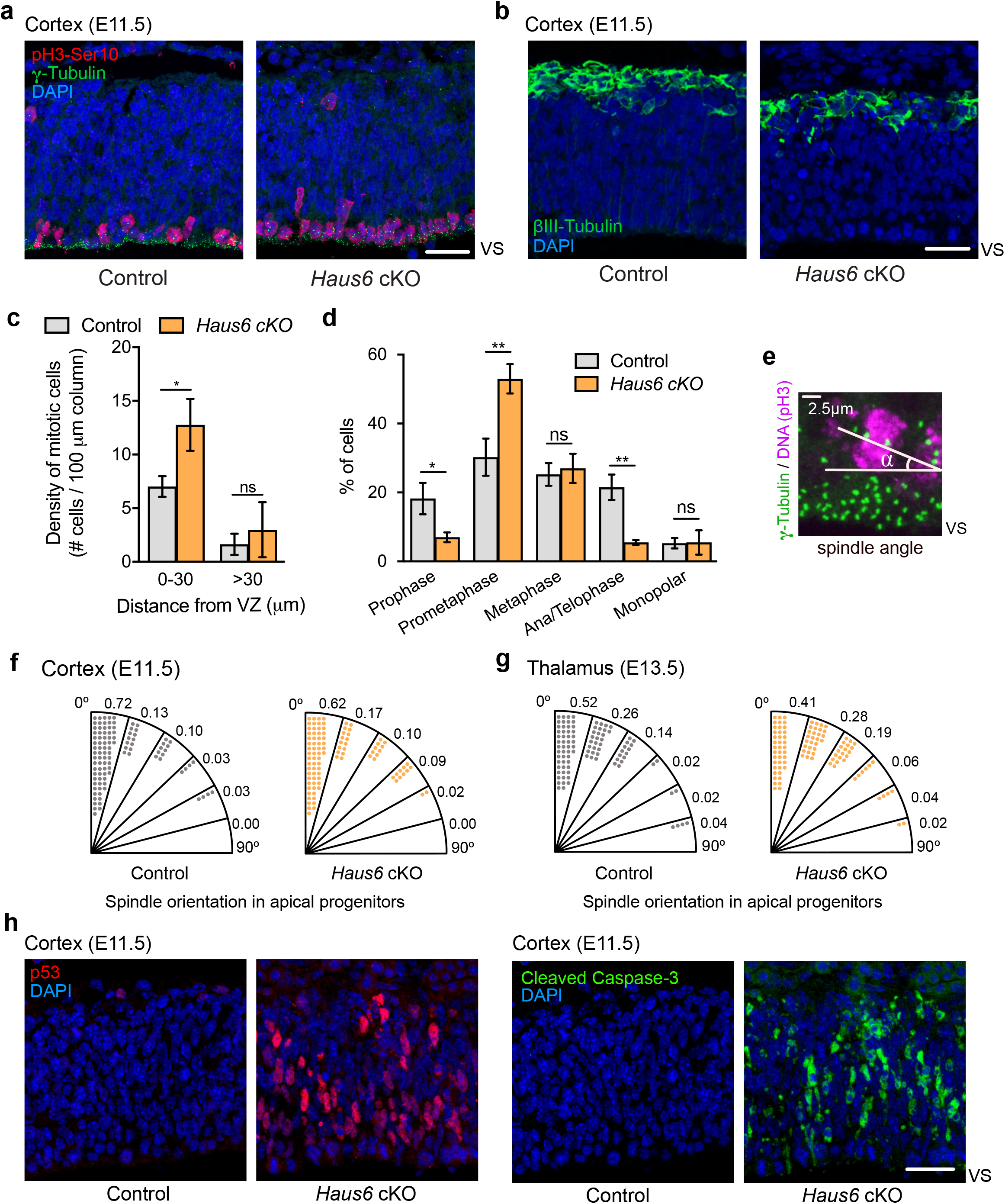
*Haus6* cKO induces mitotic defects and apoptosis at E11.5. **(a, b)** Representative images of E11.5 control (*Haus6*^wt/wt^ Nestin-Cre^+^) and *Haus6* cKO cortex (*Haus6*^fl/fl^ Nestin-Cre^+^). **(a)** Transversal brain sections were stained with antibodies against the mitotic marker phospho-Histone H3 (red), γ-tubulin (green) and DAPI to label DNA (blue). **(b)** Transversal brain sections were stained with an antibody against βIII-tubulin (green) and DAPI to label DNA (blue). **(c)** Quantification of the density of mitotic cells dividing close to the ventricular surface (<30 μm away) and in outer layers of the cortical plate (>30 μm away). **(d)** Quantification of the mitotic stages observed in mitotic cells close to the ventricular surface. **(c, d)** Plots show mean values, error bars indicate SD. n=3 Nestin-Cre control and n=2 Nestin-Cre *Haus6* cKO individual embryos. *P<0,05; **P<0,01 by two-tailed t-test. **(e)** Image illustrating how spindle orientation was quantified based on the angle formed between the spindle axis and the ventricular lining. **(f, g)** Quantification of the mitotic spindle angles in progenitors at meta/ana/telophase in **(f)** E11.5 cortex and (**g**) E13.5 thalamus. Diagrams show the distribution of the spindle angle values between 0° to 90°, grouped in 15° range intervals. Each dot represents 1% of the analyzed cells. **(f)** n=84 control and n=44 *Haus6* cKO mitotic cells from 4 and 2 individual embryos, respectively. **(g)** n=85 control and n=113 *Haus6* cKO mitotic cells from 4 and 5 individual embryos, respectively. **(h)** Representative images of E11.5 control and *Haus6* cKO cortex. Transversal sections were stained with antibodies against p53 (red) and the apoptotic marker cleaved caspase-3 (green). DNA was stained with DAPI. Scale bars: (a, b, h) 30 μm.

**Figure S3:**
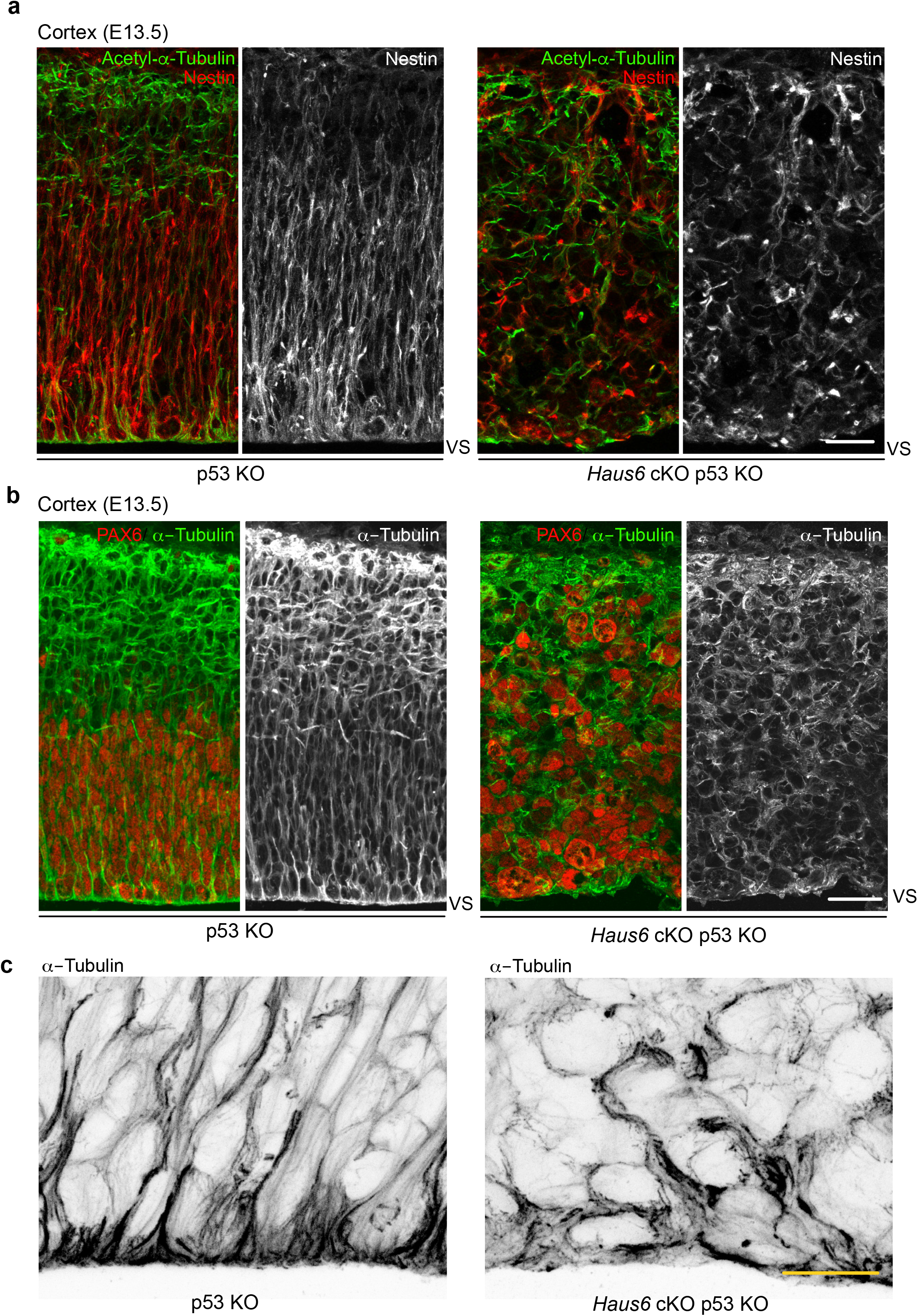
Co-deletion of *Haus6* and p53 disrupts polarity in surviving progenitors. **(a)** Representative images of E13.5 cortical sections from p53 KO control (*Haus6^fl/fl^* Nestin-Cre-p53^-/-^) and *Haus6* cKO (*Haus6^fl/fl^* Nestin-Cre^+^ p53^-/-^) embryos stained with antibodies against the apical progenitor marker nestin (red/white) and acetylated α-tubulin, a marker for stable microtubules (green). **(b)** Representative images of E13.5 cortical sections from p53 KO control (*Haus6*^fl/fl^ Nestin-Cre-p53^-/-^) and *Haus6* cKO p53 KO (*Haus6*^fl/fl^ Nestin-Cre^+^ p53-^/-^) embryos stained with antibodies against α-tubulin (green/white) and the apical progenitor marker PAX6 (red). **(c)** Magnification of the apical region of the cortex of E13.5 p53 KO control and *Haus6* KO p53 KO embryos showing microtubules stained with α-tubulin antibody. Scale bars: (a) 25 μm, (b) 35 μm, (c) 15 μm.

